# Alterations in phototransduction timing, dark adaptation and GNAT1 trafficking in *GNAT1*-edited *Xenopus laevis*

**DOI:** 10.64898/2026.10.26.761424

**Authors:** Dominic Skitsko, Beatrice M. Tam, Colette N. Chiu, Paloma Burns, Orson L. Moritz

## Abstract

GNAT1 (also called α-transducin) is the α-subunit of a heterotrimeric G-protein present in rod photoreceptors that plays a critical role in phototransduction. Photoactivated rhodopsin binds GNAT1, which stimulates phosphodiesterase to hydrolyze cGMP, closing ion channels and hyperpolarizing the rod cell.

To examine the effects of GNAT1 ablation on light-induced autophagy and light-exacerbated retinal degeneration, we previously generated genetically chimeric, *GNAT1*-edited *Xenopus laevis* using CRISPR technology. Here, we interbred these F0 animals to produce F1 animals with defined genotypes, including full knockouts, and characterized the F1 generation by Western blot, histology, and electroretinography (ERG).

In *X. laevis* tadpoles with nonsense mutations in all four *GNAT1* alleles, the GNAT1 protein was absent, and the scotopic ERG was absent, indicating full knockout. In animals with reduced GNAT1 expression, we observed increased A- and B-wave implicit times. Greater GNAT1 protein reduction was associated with greater ERG delay, demonstrating a role for GNAT1 in the timing of phototransduction.

We also identified two unexpected phenotypes linked to distinct gene edits. The Q184L- *GNAT1.L* mutation conferred an ERG phenotype similar to that of a full knockout, but with minimal reduction in GNAT1 protein levels, consistent with loss of function. This mutation is also present in the human population. A three-base-pair deletion (T183,Q184→K183) in *GNAT1.S* caused dominantly inherited diminished ERG response and defective dark adaptation, and was associated with photoreceptor structural abnormalities and GNAT1 accumulation at the connecting cilium, consistent with a dominant-negative effect on *GNAT1* trafficking.

**Summary:** Genetic disruption of *GNAT1* reveals unexpected roles in phototransduction timing, as well as loss-of-function and dominant-negative mutants that disrupt GNAT1 trafficking.

## Introduction

Rod photoreceptors are retinal neurons specialized for generating visual responses in dim light. These cells have a synaptic terminal that contacts inner retinal neurons, an organelle-containing inner segment (IS), and an outer segment (OS), a modified cilium composed of stacked membranous disks that contain the components necessary for phototransduction.

Rods generate extremely reproducible responses to signals as small as a single photon via the well-characterized phototransduction cascade. Photons activate the 7-transmembrane G-protein-coupled receptor (GPCR) rhodopsin [1] by isomerizing its covalently bound 11-*cis* retinal chromophore. Photoactivated rhodopsin proceeds through several spectroscopically distinct intermediates, including the inactive metarhodopsin I (Meta I) followed by the signaling form Meta II, with a portion of the Meta I population undergoing reversible decay into Meta III [2,3]. Meta II rhodopsin interacts with the heterotrimeric G-protein transducin, promoting GDP → GTP exchange and dissociation of the α subunit (GNAT1 aka α-transducin) from the βγ complex. Each Meta II activates 12-14 GNAT1 proteins, which in turn bind and activate cGMP-phosphodiesterase (PDE), which converts cGMP to 5’-GMP, leading to the closing of cGMP-gated cation channels, hyperpolarization of the cell, and altered synaptic vesicle release, to generate signals that are transmitted to the brain [4,5].

Thus, rhodopsin initiates the phototransduction cascade, and the heterotrimeric G protein transducin is its first amplifier. GNAT, the active signaling component, has rod- and cone-specific forms, respectively encoded by the genes *GNAT1* and *GNAT2* [6]. Mutations in human *GNAT1* cause congenital stationary night blindness (CSNB) and rare cases of rod-cone dystrophy [7,8].

Under dark-adapted conditions, heterotrimeric transducin is associated with OS membranes through the combined affinity of the three subunits’ lipid anchors [9], which are essential for OS localization [10,11]. Under lighting conditions that saturate rod signaling, activated GNAT1 is produced in excess and dissociates from OS membranes, resulting in mass translocation of GNAT1 to the IS [11–13]. This movement may allow rods to reduce their gain [14], preventing photoreceptor overstimulation and related toxicity. Additionally, GNAT1 movement to the IS may serve to regulate chromophore stores by controlling the clearance rate of Meta III rhodopsin [2].

In prolonged darkness, IS GNAT1 gradually returns to the OS via a combination of facilitated diffusion and active transport[15]. Knockout of Unc119, a GNAT1 interaction partner that solubilizes the protein within the IS, dramatically slows return of GNAT1 to the OS[16]. Return to the OS is also disrupted by interference with cytoskeletal components[17,18] and knockout of the microtubule-interacting protein Frmpd1[19].

Using genetically modified *X. laevis,* we previously observed that GNAT1 ablation via CRISPR/Cas9 gene editing suppresses the light-induced autophagy that is aggravated by the P23H rhodopsin variant [20], and increases the toxic effects of light exposure associated with the T4K rhodopsin variant [21], although these observations were made on genetically chimeric F0 CRISPR-edited animals. To assist in further study of these effects, we sought to breed these animals to generate F1 animals of defined genotype, a task complicated by the pseudo-tetraploid genetics of the *X. laevis* genome[22], which contains two *GNAT1* paralogues *GNAT1.S* and *GNAT1.L*. The majority of offspring of these F0 CRISPR-edited animals carry a combination of true *GNAT1* knockout alleles, missense and indel alleles resulting in reduced expression, and WT alleles.

Here we report that, consistent with results reported for other animals and humans, *X. laevis* tadpoles with nonsense mutations in all four *GNAT1* alleles exhibit greatly reduced scotopic ERGs and no detectable GNAT1 protein, indicating knockout. However, in animals with reduced GNAT1 expression due to missense alleles, we observed a delay in the timing of the ERG A and B waves that correlated with GNAT1 protein levels and was not explained by GNAT1’s role in signal amplification. Our results suggest a second functional role for GNAT1 in modulating phototransduction timing.

We also identified tadpoles carrying a point mutation (Q184L) that conferred an ERG phenotype similar to that of GNAT1 knockout, but with minimal reduction in GNAT1 protein. Q184L-*GNAT1.L* appeared non-functional, as it was largely confined to rod IS under dark conditions.

In a final group of animals, we observed a dominant-negative morphological and electrophysiological phenotype associated with a three-base-pair deletion in *GNAT1.S*, causing T183, Q184→K183. No photoreceptor structural abnormalities linked to GNAT1 mutations have been reported previously.

## Materials & Methods

### Generation of CRISPR/Cas9-edited and knockout *GNAT1* animals

F0 genetically edited *GNAT1* animals were previously generated by injection of sgRNA and Cas9 mRNA as described in [20,21]. The sgRNA targeted a conserved sequence GGTATCATTGAGACACAGTT present in exon 5 of both *GNAT1.S* and *GNAT1.L*. These genetically chimeric F0 animals were raised to maturity, and mated together to generate non-chimeric F1 offspring for this study. Mating/egg laying was induced by injection of 700 units (females) or 100 units (males) of human chorionic gonadotropin into the dorsal lymph sack.

### Electroretinography

Electroretinography (ERG) was performed as previously described [21,23–25]. Briefly, tadpoles were raised to Nieuwkoop and Faber stage 58 (approximately 6 weeks of age) [26] and dark-adapted overnight. Animals were anesthetized in 0.1% tricaine in 0.1X MMR until unresponsive. Tadpoles were placed on a gold ECG electrode, and a micromanipulator was used to position a silver wire electrode immersed in 1X MMR within a glass microcapillary into contact with the cornea. A lubricating drop (Genteal) was placed on the eye, and the entire preparation was placed under a Ganzfeld dome. Electrodes were connected to the headstage of a Model 1800 AC amplifier (AM Systems), and its output was connected to an Espion ERG unit (Diagnosys). The protocol consisted of 2.5, 25. 250, 750, 1250, and 2500 cd•s/m2 flashes, each repeated 5 times at 15-second intervals and averaged. The final step was a photopic flicker at 2.5 cd•s/m2 and 10 Hz over a 6500K white background. The intended use of the final flicker step was to confirm an electrical response in animals for which no scotopic ERG was detectable; however, it did not yield useful data due to rod-mediated cone suppression.

To evaluate dark adaptation, dark adapted animals were first exposed to 6 flashes of 250 cd•s/m2 blue light with 20 s between each flash to verify connection of the electrode and cornea, 5 flashes of 75 cd•s/m2 6500K white light over a 6500K white background of 250 cd/m2 with 6 s between as a bright light exposure, followed by 8 rounds of 4 flashes of 250 cd•s/m2 blue light, with 20 seconds between each round and 20 seconds between each trial.

### Euthanasia

Following ERG recording, tadpoles were immediately sacrificed by pithing. Eyes were enucleated, with the right fixed in 4% paraformaldehyde with 0.1M sodium phosphate buffer pH 7.4 and the left solubilized in 100 μL of 1:1 PBS and SDS-PAGE loading buffer with 1mM EDTA and 100μg/mL PMSF. These procedures were conducted in darkness under red light. Tail clippings were collected for DNA analysis.

### Extraction of Genomic DNA and DNA analysis

Genomic DNA was extracted from tail clippings and used as a template for PCR amplification of *GNAT1.S* and *GNAT1.L* exon 5, using DNA extraction, PCR primers, and amplification conditions identical to those of Wen et al. 2019 [20]. Amplified DNA was Sanger-sequenced (Genewiz, Azenta Life Sciences) and analyzed for mutations as described in Ghaseminijad et al. [27].

### Western blotting

We performed Western blots using solubilized eye samples as previously described [28]. 12 μL of sample was loaded per well, electrophoresed through a 10% SDS-polyacrylamide gel, and transferred to a PVDF membrane overnight. Membranes were blocked with 1% skim milk powder in PBS for 30 minutes before washing with PBST and labeling with anti-GNAT1 (1:1000, rabbit polyclonal, gift of W.C. Smith) and anti-acetylated α-tubulin (1:3000, Sigma Aldrich). After washing 3x in PBST, membranes were incubated for 2 hours with IR800-conjugated goat anti-rabbit and IR700-conjugated goat anti-mouse at 1:10,000 of 1 mg/mL solution (Rockland). Blots were washed in PBST and a final PBS wash, and imaged using a Li-Cor Odyssey imaging system. The resulting images were quantified using ImageJ [29].

### Immunolabeling and Confocal Microscopy

After 4 hours in fixative, eyes were transferred to a 20% sucrose solution for at least 30 minutes. After sucrose infiltration, eyes were embedded in OCT (Sakura Finetek), frozen, and stored at −20°C before sectioning. 12 μm cryosections were cut and labelled overnight as previously described[21], using rabbit polyclonal anti-GNAT1 (1:1000). The secondary antibody was 1:750 Cy3-conjugated donkey anti-rabbit (Jackson Immunoresearch), and counterstains were 1:100 AlexaFluor 488-conjugated wheat germ agglutinin (WGA; Life Technologies), and 1:1000 Hoechst-33342 (Sigma Aldrich). Coverslips were mounted with MOWIOL 4-88 (EMD Millipore). Labelled sections were imaged with a Zeiss 800 laser-scanning confocal microscope or a Zeiss 880 Airyscan microscope and Zen software, using a 40X water-immersion NA 1.2 objective. Maximum-intensity projected images were generated using Zen software from optical sections with a consistent spacing of 0.3 microns per section. Within each experiment, all images were acquired with the same gain and contrast settings. For experiments involving quantification of confocal micrographs, comparisons were made only between images acquired under the same settings.

### Generation of microscopy figures

16-bit images acquired in Zen software were post-processed and assembled into figures using Adobe Photoshop. Hoechst and WGA counterstain channels were adjusted non-linearly to improve image clarity, while GNAT1 labelling was adjusted linearly.

### Optical coherence tomography retinal imaging

Post-metamorphic frogs were anesthetized in 0.1% buffered tricaine. When the animals no longer responded to touch, retinal imaging was performed with a Phoenix Micron IV camera and an OCT system designed to image mouse eyes.

## Results

To generate *X. laevis GNAT1.S* and *GNAT1.L* double knockouts, we began by interbreeding genetically chimeric CRISPR-modified F0 animals previously generated in our lab [20,21]. F1 tadpoles generated by mating of our selected candidate F0 animals were raised to 6 weeks of age before an initial screening for knockouts by electroretinography. The use of genetically chimeric F0 animals and the presence of four *GNAT1* alleles meant that alongside the desired double knockouts, we also generated *X. laevis* with incomplete *GNAT1* knockout, due to genotypes that included some WT alleles and/or edited alleles that result in reduced expression. Thus, we obtained a greater variety of expression levels than would be achievable in diploid animals such as mice, allowing us to correlate the effects of reduced *GNAT1* expression with properties of the ERG.

### GNAT1-knockout X. laevis

As predicted, we identified a subset of animals with absent rod ERGs (Fig. 1A/B). Following electroretinography, we euthanized a subset of these animals; one eye was solubilized for Western blot analysis, another was fixed for microscopy, and genomic DNA was isolated for genotyping. Immunohistochemical analysis confirmed the absence of GNAT1 labeling in both rod IS and OS (Fig. 1C). Absence of GNAT1 was also confirmed by Western blot (Fig. 1D). Sanger sequencing identified frameshift mutations generating premature stop codons (nonsense mutations) in all four *GNAT1* alleles, confirming successful generation of *GNAT1* knockout *X. laevis* (not shown). Consistent with previous results in *X. laevis* and other species [20,21,30] we did not observe any retinal degeneration in these animals. Adult knockout animals imaged by OCT presented with normal OS thickness and morphology, as did animals with the additional specific edits of interest reported below (Fig. 1E; n=3 WT OS 38.02±4.96 μm, n=9 KO OS 35.06±5.18 μm).

**Figure 1.**
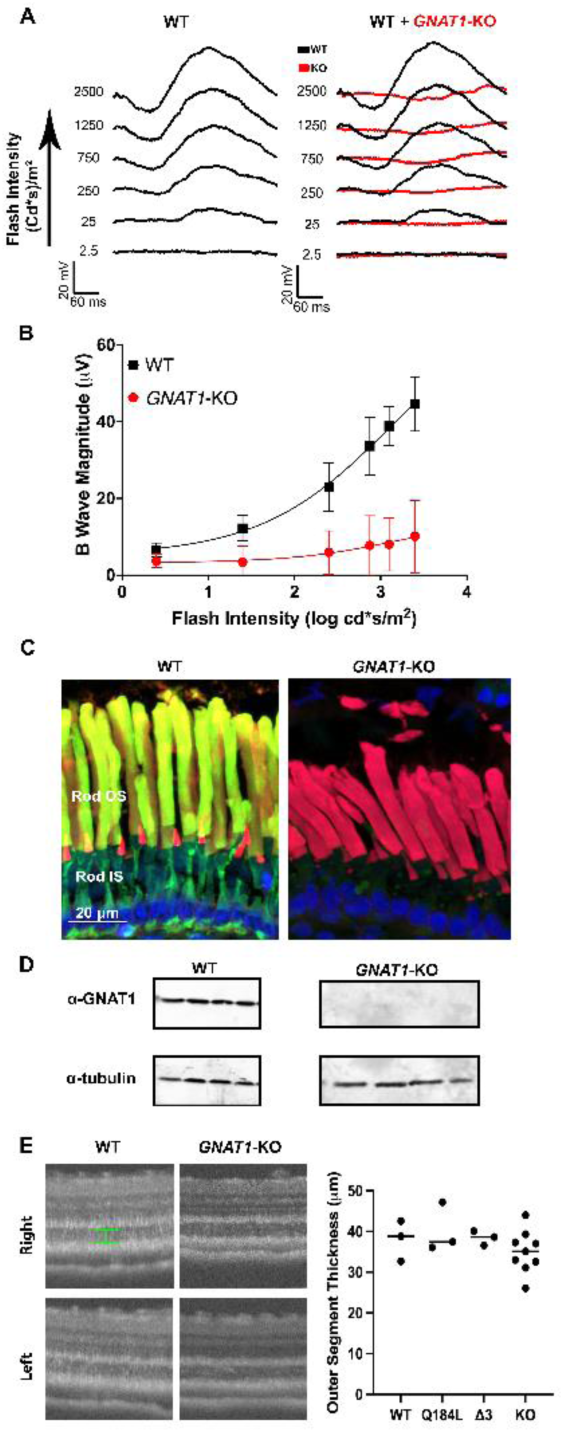
Identification of *GNAT1*-knock out *X. laevis:* **A, B:** Potential *GNAT1*-knockout animals were initially identified by electroretinography screening, where they presented with absent scotopic ERGs relative to WT animals. Traces shown are an average of n=6 WT (black), n=8 KO (red). Reduction in B wave response amplitude was statistically significant by two-way ANOVA (B; p<0.0001 for all factors). **C:** Suspected knockouts were confirmed immunohistochemically via GNAT1-labeling (green), which was absent in knockouts (right) relative to WT (left). Red = wheat germ agglutinin, blue = Hoechst 33342. **D:** Absence of GNAT1 was further confirmed by Western blotting of solubilized eyes. Acetylated α-tubulin labelling served as a loading control. **E:** Adult (>6 months) knockout retinas, as well as retinas with other edits characterized here, all presented with normal appearance via OCT (OS thickness measured between the marked green lines).

### Reduction of *GNAT1* Expression Correlates with Increases in ERG Implicit Time

In addition to the expected absent ERG phenotype associated with *GNAT1* knockout, we also noted a number of F1 offspring with an unexpected ERG phenotype consisting of a variable reduction in ERG amplitude and a significant delay in the A and B wave implicit times, with some animals exhibiting responses that were significantly delayed at all flash intensities (Fig 2C). These changes were most apparent at a flash intensity of 2500 Cd•s/m^2^, where the average WT A-wave implicit time was 69±9.1 ms (n=10) while partial *GNAT1-*KO implicit times ranged from 62 ms to 196 ms. The average WT B wave implicit time was 178±13.4 ms (n=10), while partial knockouts ranged from 176 ms to 386 ms. When the implicit time of the ERG was quantified and plotted against GNAT1 expression values derived from Western blots, a clear relationship emerged (Fig. 2D). The relationship between A wave implicit time and expression level fit a first order exponential decay relationship with an R^2^ of 0.82 and a *GNAT1* expression of 6.5% producing a half-maximal effect, while the relationship between B wave and *GNAT1* expression level fit a linear relationship with an R^2^ of 0.57.

**Figure 2.**
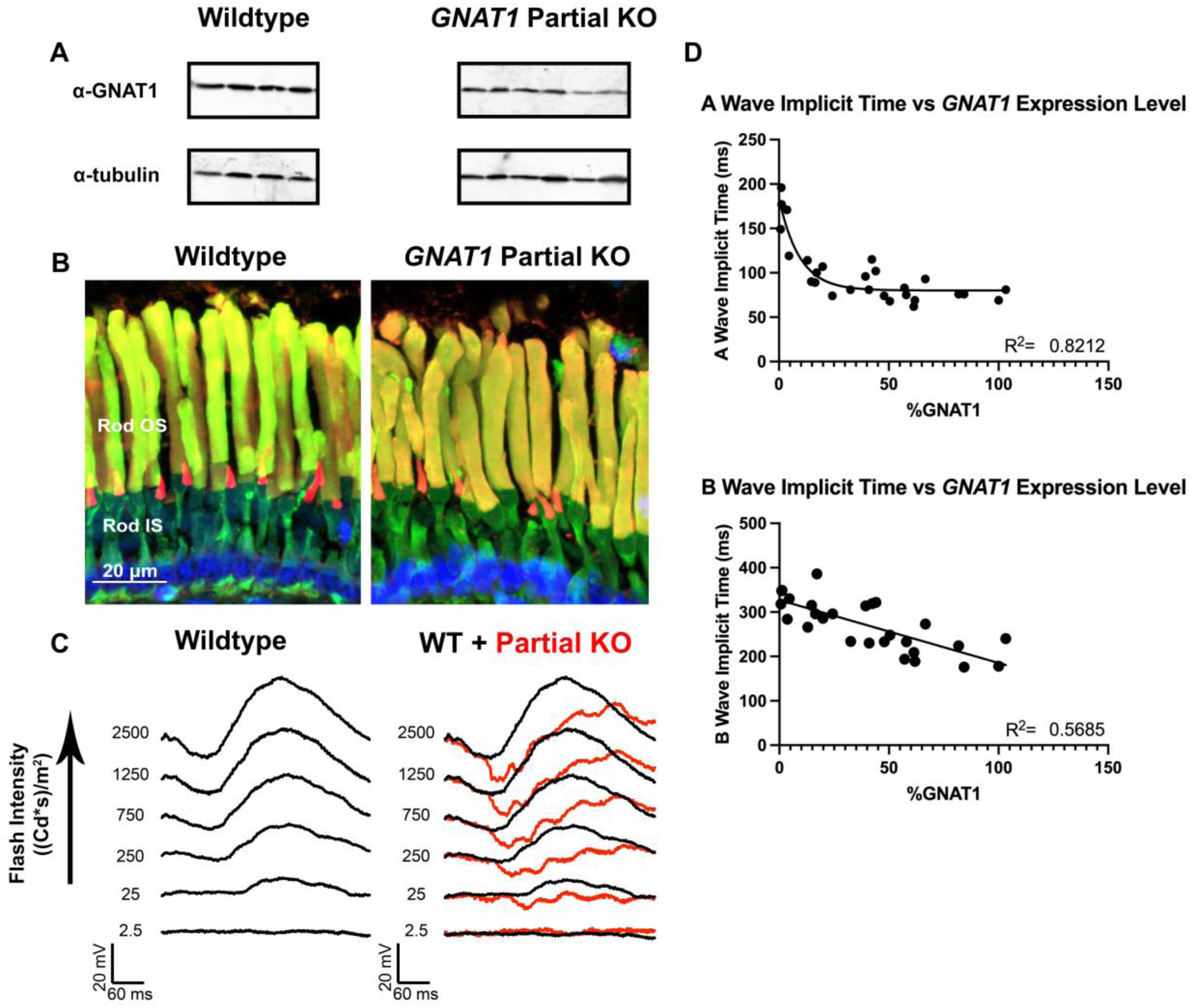
Reduced GNAT1 expression correlates with a delayed ERG. **A:** Many edited animals were not full knockouts and showed reduced expression of GNAT1 by western blot. **B:** Similarly, confocal microscopy frequently showed reduced GNAT1 labeling intensity relative to WT. Green = GNAT1, Red = wheat germ agglutinin, blue = Hoechst 33342 **C:** Animals with altered (delayed) ERG responses were also noted. (average of n=6 WT in black, and a representative delayed trace superimposed in red). **D**. Implicit times of ERG A- and B-waves were plotted against GNAT1 expression levels obtained from Western blots. The relationship between expression level and A-wave implicit time fits a first-order exponential decay (R^2^ = 0.83), **with a** half-maximal effect at 6.5% expression. The relationship of B-wave and expression is linear (R^2^ of 0.57). Animals without discernible A- and B-waves (probable knockouts) were excluded from the analysis.

### Functional Knockout of *GNAT1* Associated with a One Base Pair Substitution in *GNAT1.L*

While conducting the ERG and Western blot screening described above, we identified a subset of animals with an absent ERG (Figure 3A) but abundant GNAT1 labelling, in which GNAT1 was mislocalized to the RIS under dark conditions (Figure 3B). Western blots demonstrated that, unlike other animals lacking an ERG (Figure 1A-D), GNAT1 protein was relatively abundant (Figure 3C), indicating that these animals were functional *GNAT1* knockouts. Sanger sequencing identified a one-base-pair substitution causing the point mutation Q184L in *GNAT1.L* in all *X. laevis* with this “functional KO” (high expression, absent ERG) phenotype. In our initial analysis of 12 tadpoles derived from a single mating, all other alleles present in all 12 animals were either frameshift mutations in *GNAT1.L* or *GNAT1.S* consistent with knockout, or deletions in *GNAT1.S* that were also present in animals that lacked this phenotype, clearly implicating the Q184L mutation in the phenotype. (n=3 out of n=12 total, P = 0.000532 by Chi-square test). In subsequent experiments, when Q184L was observed in combination with wild-type alleles, a robust ERG was present, indicating recessive inheritance of the ERG phenotype. Adults presented with normal retinal morphology via OCT (Fig. 1E, n=3 WT OS 38.02±4.96 μm, n=3 Q184L OS 40.22±6.05 μm)

**Figure 3.**
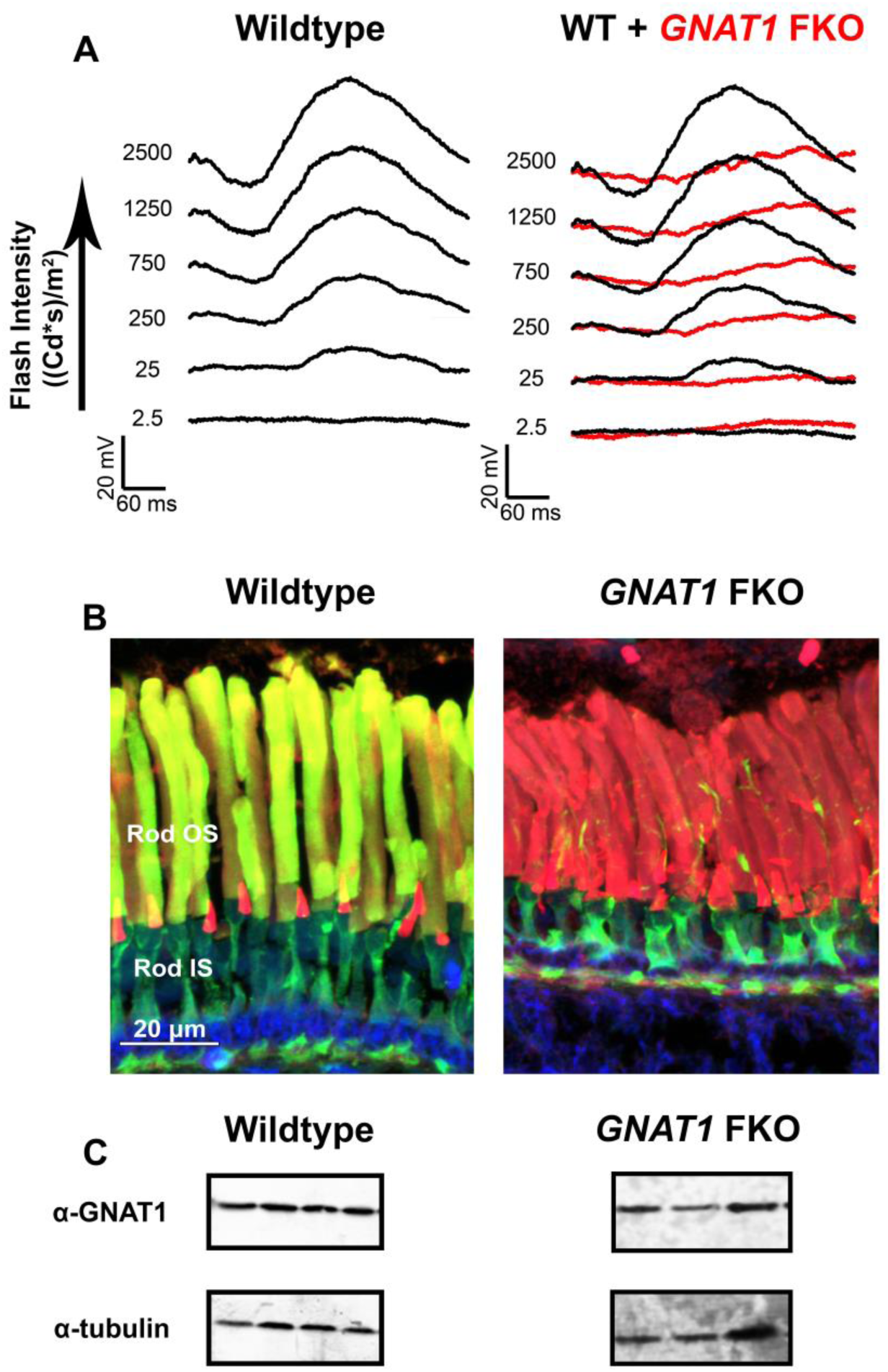
Identification of *GNAT1* functional knock-out animals. **A, B:** We identified a group of animals that resembled *GNAT1* knockouts due to their absent ERG (average of n=6 WT in black, average n=3 functional knockouts in red), but with abundant GNAT1 labeling (Green = GNAT1, Red = wheat germ agglutinin, blue = Hoechst 33342) confined to the rod IS and the ciliary axoneme. **C:** Western blotting confirmed the presence of GNAT1 within these samples.

**Figure 4.**
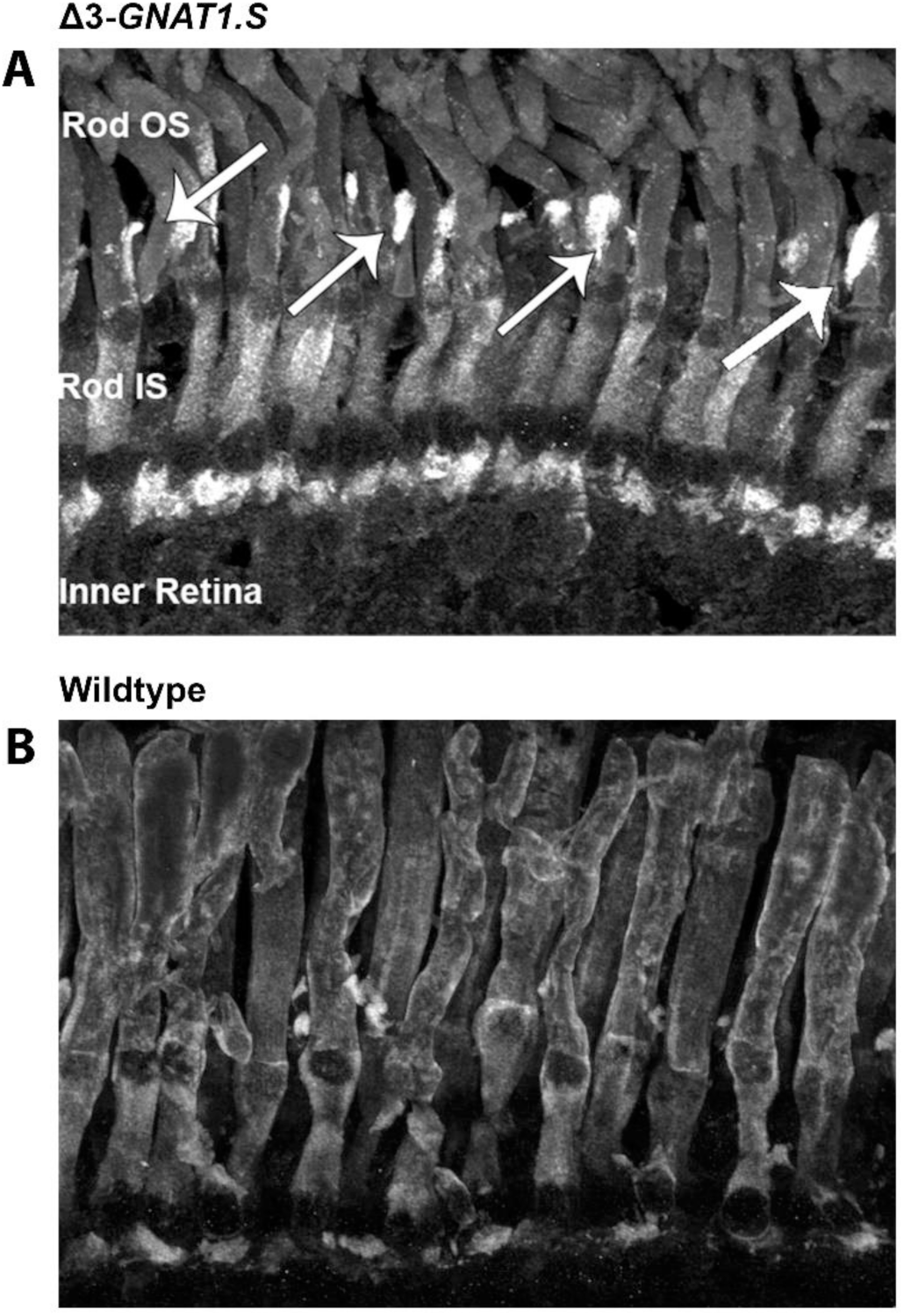
Identification of animals with GNAT1-postive abnormal protrusions. **A:** Examination of samples generated from the mating of F0 animals via Airyscan super-resolution microscopy revealed a group of *X. laevis* tadpoles with GNAT1 confined to the rod IS and abnormal morphological structures adjacent to the OS. These structures were not present in WT animals, in which GNAT1 localized to the OS (B).

### *X. laevis* carrying a Δ3 indel in *GNAT1.S* have abnormal rod morphology

We also identified a second subset of animals based on their abnormal ERG and histological appearance, which presented with abnormal GNAT1-positive structures at the base of the rod OS (Fig. 7A). In these animals, GNAT1 label was restricted to the IS and these structures. Genotyping linked this phenotype to a three base-pair deletion in *GNAT1.S*, causing the mutation T183-Q184 → K183 (abbreviated Δ3-*GNAT1.S* in the following text and figures). The morphological phenotype was apparent in 5 animals expressing Δ3-*GNAT1.S*, and all other mutations present in these animals were also observed in animals lacking this distinct phenotype (n=5 with phenotype out of n=11 total, P = 0.000911 by Chi-square test). All animals with the observed phenotype also had at least one wildtype *GNAT* allele, suggesting dominant inheritance. Adult F1 animals imaged via OCT presented with normal retinal layer thickness (Fig. 1E; n=3 WT OS 38.02±4.96 μm, n=3 Δ3-*GNAT1.S* OS 38.42±1.76 μm).

### The Δ3-*GNAT1.S* phenotype is dominantly inherited

We raised a number of these animals to maturity and subsequently bred an F1 animal carrying the Δ3-*GNAT1.S* mutation with a WT animal; these F2 offspring were analyzed as described above. The F1 animal (and therefore 50% of its offspring) also carried a 12-bp deletion in *GNAT1.S* that was not associated with the morphological phenotype. F2 offspring carrying the Δ3-*GNAT1.S* mutation developed the morphological phenotype observed in F1 animals, with GNAT1 labelling strongly confined to the IS and the abnormal GNAT1-positive protrusions (n= 5; Fig. 8C). These animals also had a significantly reduced ERG compared to WT (Fig. 8D). In contrast, the siblings of these animals carrying the 12-base pair deletion in *GNAT1.S* presented with minimal ERG alteration and GNAT1 present in the OS, with somewhat more GNAT1 labeling in the IS than typically seen in WT animals (n=6; Fig. 8A/B). This was consistent with misfolding of a fraction of the total GNAT1 induced by the 12-bp deletion. Thus, the phenotype produced by Δ3-*GNAT1.S* is dominant and distinct from that produced by other *GNAT1.S* mutations.

### GNAT1 is present in the outer segments of Δ3-*GNAT1.S-*expressing rods after exposure to steady light

We wondered whether the build-up of GNAT1 in the abnormal structures in the vicinity of the connecting cilium could be related to abnormal light-induced GNAT1 trafficking. Therefore, we exposed overnight dark-adapted siblings of the animals described above to 10 minutes of constant 1700 lux light, sacrificed a group immediately, then returned them to darkness and sacrificed them at intervals of 4, 32, 60, and 120 minutes. We observed GNAT1 labelling in the OS of all animals (n=15), both those expressing Δ3-*GNAT1.S* and those expressing the alternate Δ12-*GNAT1.S* mutation, under dark conditions and during dark adaptation (Fig. 9). The unusual GNAT1-positive protrusions were not present.

### The photopic flicker of the ERG protocol is required to produce the altered labelling pattern and morphology in Δ3-*GNAT1.S* animals

After determining that steady bright light was insufficient to produce the Δ3-*GNAT1.S* phenotype, we investigated whether specific aspects of the ERG protocol were critical for inducing the phenotype by exposing sets of animals to the entire protocol, or specific portions of the protocol. Our phenotype was produced by combinations of steps containing a photopic flicker stimulus that was the final step of our protocol (n=15) including the photopic flicker alone (n=5, Fig. 9), indicating that the photopic flicker step was the stimulus that induced the Δ3-*GNAT1.S* morphological phenotype. Notably, the cumulative light exposure of the flicker step (1500 cd/m^2^) was considerably less than the 10-minute light exposure used in Fig. 9 (170,000 cd/m^2^), suggesting that it was not total light exposure but the flickering quality of the light that was essential (Table 2).

### Return of GNAT1 to the OS Following a Bleaching Stimulus is Delayed in Δ3-*GNAT1.S* Expressing Animals

We investigated the apparent involvement of light-induced GNAT1 trafficking in our Δ3-*GNAT1.S* phenotype using offspring generated by crossing our Δ3-*GNAT1.S*-F1 animal with wildtype. We dark-adapted tadpoles overnight, exposed them to our translocation-inducing lighting regimen, then returned them to the dark. We euthanized and collected eyes from overnight dark-adapted, light-adapted, and 4/32/60/120-minute dark-adapted animals (n=4 per time point), labeling them as described above and quantifying the label using ImageJ. Comparing changes in the ratio of the mean GNAT1 signal intensity in the OS and IS for wildtype and Δ3-*GNAT1.S* animals revealed that the rate of return and endpoint were dramatically reduced in Δ3-*GNAT1.S* retinas (Fig. 6A), indicative of obstructed OS trafficking of GNAT1. These differences were highly significant by two-way ANOVA (n=10, 5 rods from two animals each per treatment; P interaction = 1.03•10^-11^, P time = 3.06•10^-11^, P genotype = 9.19•10^-14^).

In order to confirm that our observed changes in OS/IS ratio were due to protein movement and not consistently high IS expression of GNAT1 in the Δ3-*GNAT1.S* group, we compared the mean signal intensity for each compartment under fully dark and light-adapted conditions. Light-adapted animals presented with increased GNAT1 within the IS and decreased GNAT1 signal in the OS for both groups (Fig. 6B/C), and all light-induced changes within genotypes were significant by two-way ANOVA. This result confirmed that the altered GNAT1 signal ratios in Fig. 6A were due to light-induced translocation of GNAT1 between OS and IS, with the decreased rate of recovery and endpoint in Δ3-*GNAT1.S* animals reflective of impaired translocation of GNAT1 to the OS.

### Dark Adaptation Is Reduced in Δ3-*GNAT1.S* Expressing Animals

After confirming that our Δ3-*GNAT1.S* mutation has a dominant-negative effect on GNAT1 trafficking during dark adaptation, we examined the electrophysiological consequences of this defect. We exposed dark-adapted animals to bright light, followed by regularly spaced low-intensity flashes in darkness. As expected, WT responses to the flashes increased in magnitude progressively as the rods dark-adapted (Fig. 7A). In contrast, although light-adapted Δ3-*GNAT1.S* (n=10) animals displayed a similar initial response magnitude to WT controls (n=5), their B wave magnitude did not significantly recover over the course of the protocol (Fig. 7A/C). Comparing WT and Δ3-*GNAT1.S,* and Δ12-*GNAT1.S* (n=4) animals revealed a highly statistically significant interaction between genotype and time via two-way ANOVA (P interaction = 4.54•10^-5^, P time = 3.17•10^-6^, P genotype = 5.34•10^-12^; Figure 7). Tukey’s post-hoc comparison revealed no significant differences at t=0 or t=100. Δ12-*GNAT1.S* was not significantly different from wildtype until t=400 (p=1.52•10^-2^), while Δ3-*GNAT1.S* significantly differed from both groups at t=200 (P wildtype = 4.66•10^-5^, P Δ12-*GNAT1.S* = 7.31•10^-3^), t=300 (P wildtype = 5.00•10^-9^, P Δ12-*GNAT1.S* = 1.72•10^-2^), and t=400 (P wildtype = 4.66•10^-5^, P Δ12-*GNAT1.S* = 7.31•10^-3^). These results confirm that Δ3-*GNAT1.S* exhibits a significant effect on dark adaptation distinct from deficits caused by other *GNAT1* mutations.

## Discussion

We have successfully generated, identified, and characterized *GNAT1-*KO and *GNAT1*-edited *Xenopus laevis.* The *GNAT1*-KO phenotype was consistent with previous reports from mice and humans[8,30], presenting with an absent scotopic ERG, no GNAT1 present by Western blot, no labeling of GNAT1 via immunofluorescence microscopy, frame-shift mutations in all *GNAT1* alleles, and no apparent retinal degeneration. Residual responses at high flash intensities were similar to responses from RPE65-KO and F0 GNAT1-KO *X. laevis*[21,23]. The residual responses could be due to the activity of other photosensitive retinal cell types, such as intrinsically photosensitive retinal ganglion cells. These *GNAT1* knockout animals will provide a useful background for future investigation of the role of phototransduction in retinal degenerations caused by rhodopsin mutations by allowing us to express disease-causing transgenes on the *GNAT1*-KO background.

In the process of generating these knockout animals we obtained several unexpected results that provide insight into the function of GNAT1. Firstly, we found that reductions in GNAT1 expression level below 50% result in a delay of the ERG. Interestingly, there is precedent in the literature for low GNAT1 expression altering the kinetics of phototransduction. In their 2019 paper, Yue et al. [31] expressed wild-type GNAT1 at ∼6% of normal levels on a *GNAT1*-KO background, a value remarkably close to our observed GNAT1 expression level that induces a half-maximal effect (6.5%). These mice exhibited a ∼1.5x increase in integration time, time to peak, and recovery time constant. This range agrees with our observed ∼1.75x increase in A wave implicit time at 6% GNAT1 expression. The presence of this delay when WT GNAT1 is expressed at low levels [31] and with a variety of gene edits (this study) suggests that this effect is directly related to decreased GNAT1 expression rather than to a mutant protein with altered function.

The fact that a reduction in GNAT1 levels alters the timing of the ERG suggests a second functional role for GNAT1, in addition to its role in amplifying the phototransduction cascade. In fact, previous *in vitro* work demonstrated that in isolated discs, the presence of GNAT1 results in the production of excess active Meta II rhodopsin compared to the intermediate Meta I form [32]. Additionally, a 2004 paper from Zimmerman et al. demonstrated that, *in vitro*, the rate of conversion of the retinal storage photointermediate Meta III rhodopsin into the active signaling Meta II rhodopsin form is dependent on the concentration of GNAT1, with evidence for a direct interaction between GNAT1 and Meta III [2].

Therefore, the delayed ERG response we observe may reflect these effects *in vivo*. With decreased expression of GNAT1, conversion of the initially produced Meta I to the signaling Meta II form could be delayed, similarly delaying the ERG A-wave. Additionally, any Meta III generated may not be efficiently converted to Meta II, amplifying this effect. As these intermediates gradually convert to Meta II, the photoresponse would be prolonged, resulting in an overall delayed ERG. This effect only becomes prominent at very low expression levels, as seen in our observed half-maximal effect at 6.5% expression, and the transgenic mouse of Yue et al. [31] expressing at ∼6%, explaining why it is not observed in heterozygous *GNAT1*-KO mice in which only a 25% reduction in expression occurs[30].

The existence of a direct relationship between GNAT1 and rhodopsin intermediates raises interesting questions regarding the role of GNAT1 in light-induced retinal degeneration. Notably, previous work from our lab implicates toxic photointermediates in T4K rhodopsin-induced retinal degeneration, with CRISPR-based ablation of *GNAT1* dramatically exacerbating this effect[21]. GNAT1 may interact with toxic intermediates produced by mutant rhodopsins upon activation, in a manner similar to MetaI and MetaIII, thereby facilitating their conversion into Meta II and subsequent decay to rod opsin and free chromophore.

Alternatively, the ERG delay we observed could be related to the organization of rhodopsin and transducin complexes within OS disks. Cryoelectron microscopy analysis reveals rhodopsin is organized into tracks of dimers[33]. Among other possible reasons for this structure, a role in phototransduction kinetics has been proposed, with reproducibility of single-photon responses promoted by the preassembly of a relatively consistent number of transducin complexes with each rhodopsin track. In this scenario, large decreases in α-transducin expression would delay phototransduction, as some tracks would have no pre-complexed transducin, and activated rhodopsin would need to recruit α-transducin from neighbouring tracks, a slower process[33]. Twelve to fourteen GNAT1 are activated per active rhodopsin[31]. With an IC50 of 6.5% for the relationship between *GNAT1* expression and A wave timing, this means that the greatest impacts on phototransduction in our model are seen when, on average, less than one GNAT1 is activated per rhodopsin, agreeing with predictions made by Gunkel et al.[33].

In addition to the general effects of reduced GNAT1 expression on the ERG, we also characterized the effects of two distinct *GNAT1* mutations. In the first case, we identified a *GNAT1.L*-Q184L mutation that abolishes GNAT1 function while maintaining robust GNAT1 expression, i.e., a functional knockout. By a remarkable coincidence, a mutation equivalent to *GNAT1.L-*Q184L has been linked to a silencing phenotype in the yeast G protein α-subunit analogue *GPA1*[34,35]. This mutation suppresses the effects of a constitutively active βγ mutant by altering interactions between the α and βγ subunits to increase affinity. Q184L is located between the switch I and II regions of the protein, which change conformation in response to GTP binding and dissociation from βγ-transducin [36]. Therefore, it is likely that this mutation induces a conformational change that specifically alters binding of α-transducin to βγ, preventing transport to the OS without causing biosynthetic defects. This proposed mechanism also aligns with the recessive nature of the phenotype in our model; the Q184L mutation does not affect the entire GNAT1 population of the cell and is compensated for by expression from other *GNAT1* alleles. In an additional remarkable coincidence, we searched the GNOMAD database and found that an equivalent Q184L *GNAT1* variant is present as a rare allele in the human population [37]. Although it is not associated with any reported disease cases, our results suggest it is likely a pathogenic variant capable of causing autosomal recessive CSNB.

Finally, we identified a dominant-negative *GNAT1* variant, caused by a 3bp deletion, T183-Q184 → K183 associated with ERG and dark adaptation deficits and a light-induced alteration in rod morphology. This phenotype is related to deficits in GNAT1 trafficking to the OS rather than a misfolding-induced mislocalization. The morphological defect associated with Δ3-*GNAT1.S* mutant occurs only under specific lighting conditions, with GNAT1 localized to the OS when dark-adapted (Figure 5). Following light adaptation, GNAT1 returns to the OS in both groups, but to a lesser extent in Δ3-*GNAT1.S X. laevis* than in WT (Figure 6). We also observed deficits in dark adaptation in these animals (Figure 7), consistent with a phenotype resembling human CSNB.

**Figure 5.**
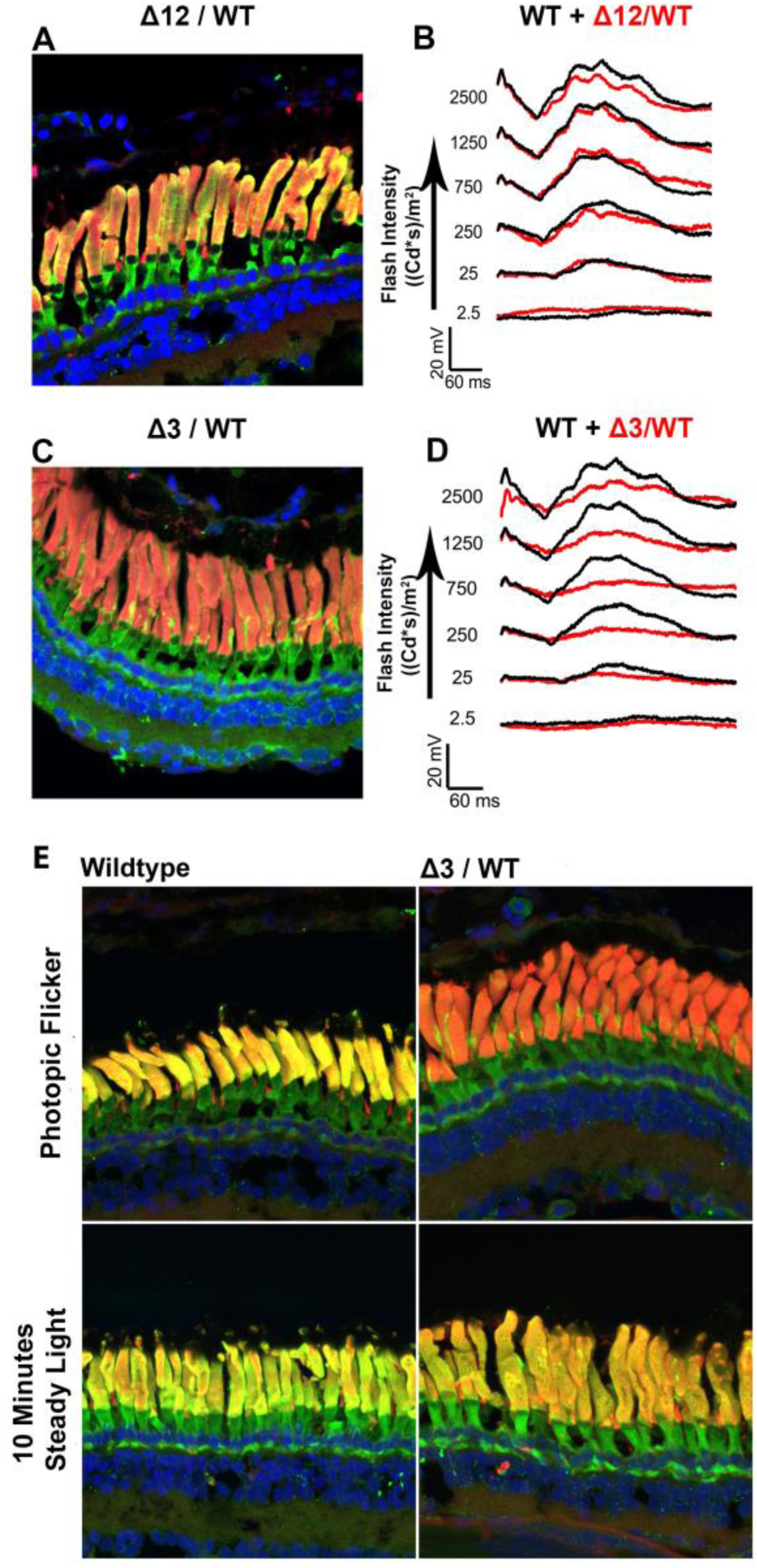
Dominant phenotypes associated with Δ3-*GNAT1.S*. **A-D**: F2 heterozygous Δ3-*GNAT1.S* x WT animals have IS localization of GNAT1 with protrusions and a reduced ERG (D; average of n=5 WT, n=5 Δ3)., while siblings carrying the alternative Δ12-*GNAT1.S* mutation presented with some IS GNAT1 signal and a normal ERG (**B**; average of n=5 WT, n=5 Δ12) (Green = GNAT1, Red = wheat germ agglutinin, blue = Hoechst 33342). **E:** The abnormal localization of GNAT1 in Δ3-*GNAT1.S* animals was dependent on exposure to the flicker step of the ERG protocol and did not occur in WT animals or Δ3-*GNAT1.S* animals exposed to a steady light source.

**Figure 6.**
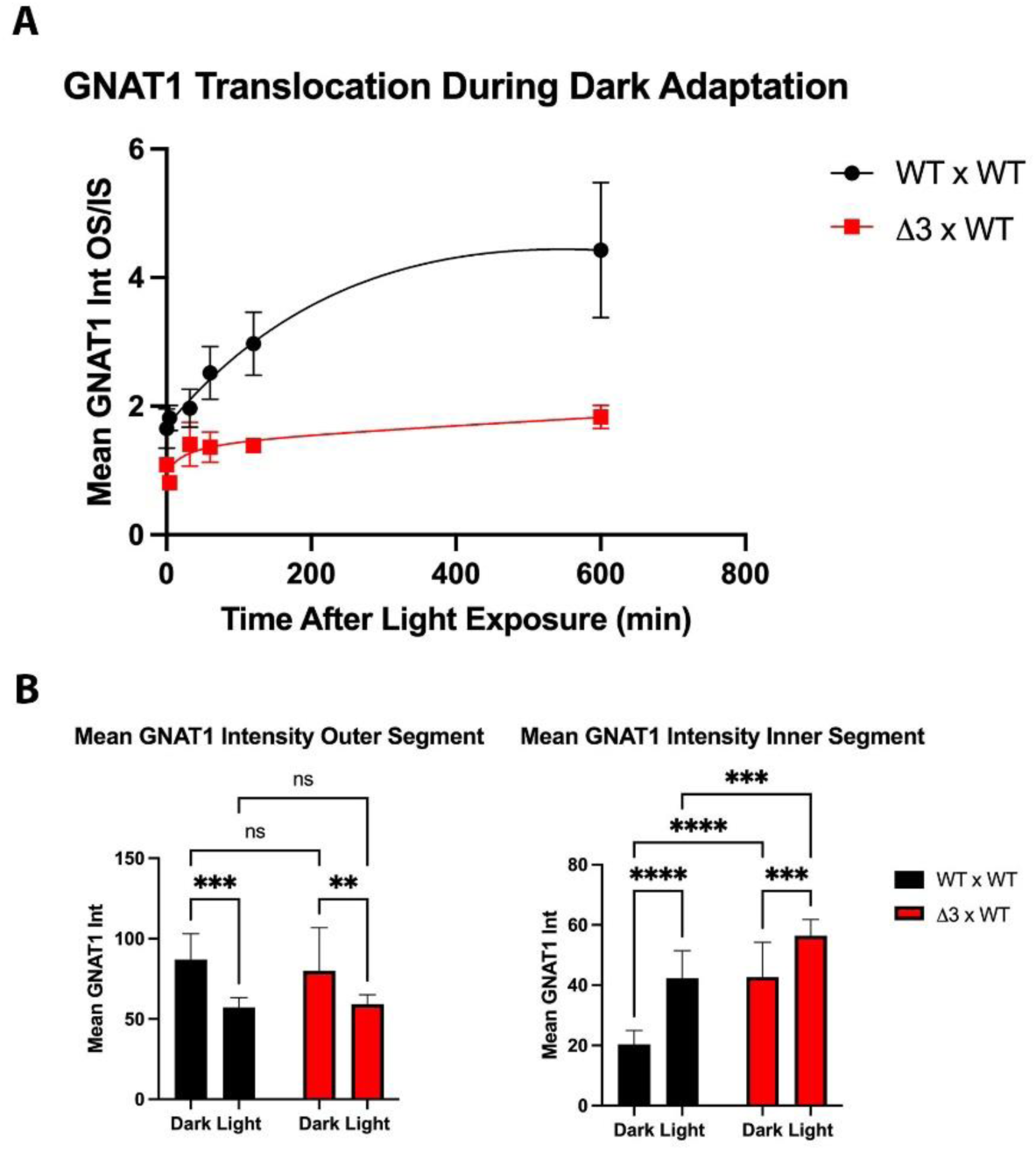
GNAT1 return to the outer segment is significantly altered in Δ3 x WT-*GNAT1.S* animals. **A:** ratio of the mean GNAT1 signal intensity for rod OS and IS at increasing time points following light exposure (n=5 rods from two animals for each time point) shows that the return of GNAT1 to the OS during dark adaptation is reduced in Δ3 x WT-*GNAT1.S* animals. (Two-way ANOVA time vs genotype, p<0.0001 for all factors). B: Comparison of the mean signal intensity of rod OS (left) and IS (right) shows that changes in the GNAT1 levels occur in both IS and OS (WT/WT OS p=0.0002, IS p<0.0001; Δ3/WT OS p=0.0071, IS p=0.0006), confirming GNAT1 translocation occurs in Δ3/WT-*GNAT1.S* rods, albeit altered compared to WT. Mean IS intensity is elevated vs WT in both light and dark-adapted conditions (dark p<0.0001, light p=0.0004) while no significant difference in the OS was observed (dark p=0.3432, light p=0.7853). *= p<0.05, **= p<0.01, ***= p<0.001, ****= p<0.0001.

**Figure 7.**
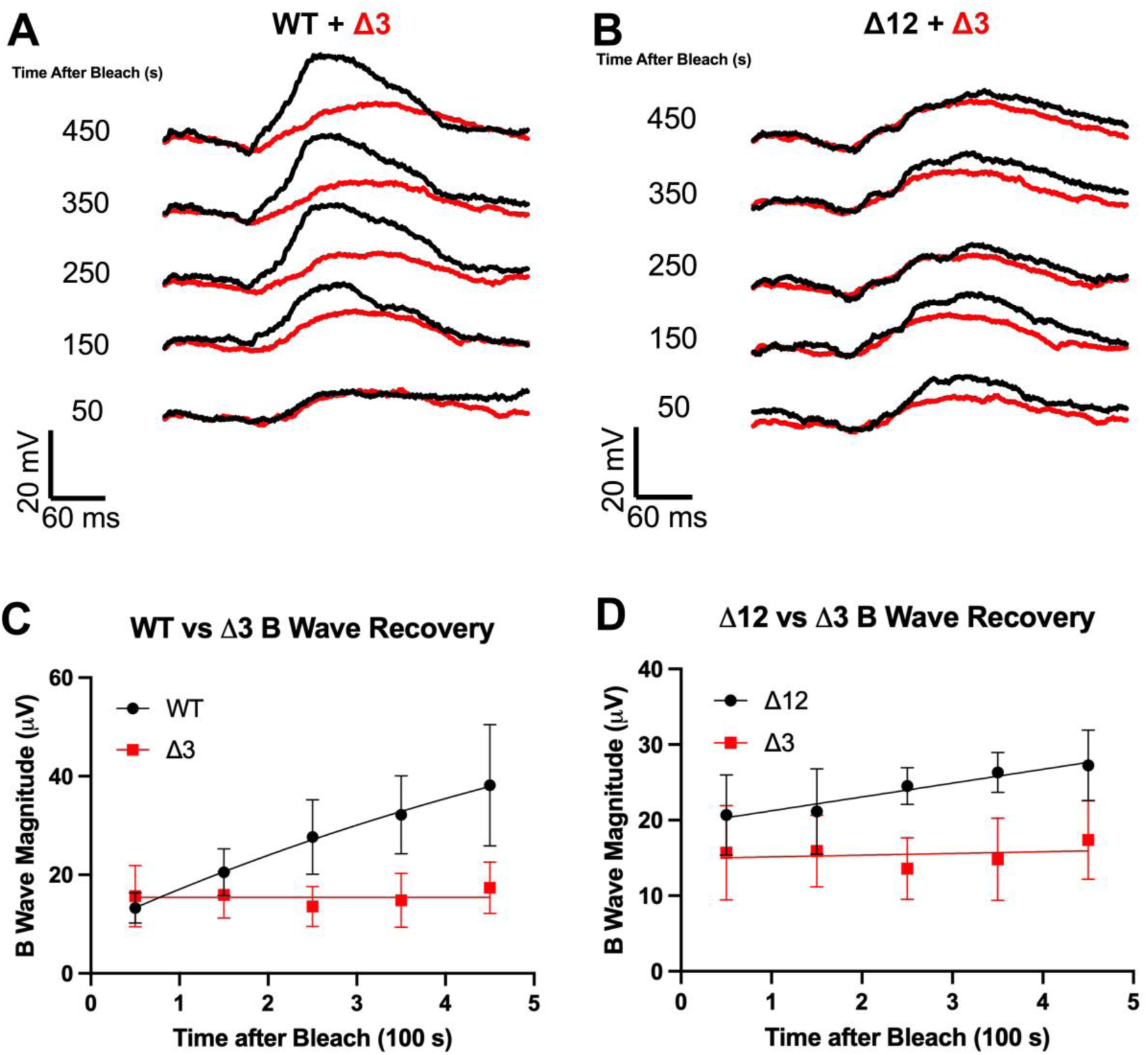
Delayed dark adaptation in Δ3 x WT-*GNAT1.S* animals. A, C: ERG recordings of responses to repeated low-intensity light flashes following a bright light stimulus show that dark adaptation of Δ3 x WT-*GNAT1.S* animals (n=10) is significantly impaired when compared to wildtype (n=5). B, D: A similar result was obtained when the same Δ3/WT-*GNAT1.S* animals were compared to their Δ12/WT-*GNAT1.S* siblings (n=5).

T183-Q184 → K183, the three-base-pair deletion producing structural and electrophysiological abnormalities, occurs in a region adjacent to the switch II region that is known to stabilize interactions between the N-terminal myristylation site with UNC119 [16,38]. As UNC119 is essential for GNAT1 trafficking to the OS, issues with association between GNAT1 and UNC119, such as delayed release of GNAT1, may be the cause of the structural abnormalities observed; for example, Δ3-*GNAT1.S* could block the transport of co-expressed WT GNAT1 by saturating binding to UNC119. The light-induced protrusions containing high concentrations of GNAT1 may represent a backlog of GNAT1 trafficking at the site of UNC119 binding. The repeated photopic flicker flashes may exaggerate the phenotype by producing closely spaced rod-saturating stimuli, driving multiple waves of GNAT1 translocation that exceed those induced by a non-flickering stimulus.

The trafficking defects we observed could also be caused by interference with the Gpsm2/Frmpd1 active transport pathway. Knockout of Frmpd1 results in decreased return of GNAT1 to the OS and deficient dark adaptation measured via ERG [19], similar to that we observed with the Δ3-*GNAT1.S* mutant. Gpsm2 is present at a ∼1:15 ratio compared to GNAT1 [39], so saturation could occur with only one mutant allele. Unfortunately, the specific region of the GNAT1 protein interacting with Gpsm2 and FRMPD1 remains unclear.

Regardless of the underlying mechanism, the Δ3-*GNAT1.S* mutation provides an interesting model for examining light-induced GNAT1 movement, its return to the OS and its role in cellular health and dark adaptation. Our observed phenotype could share an underlying mechanism with extremely rare dominantly inherited *GNAT1*-related disorders, including dominant congenital stationary night blindness[40] and rod-cone dystrophy. Moreover, this mutant may also allow us to examine the role of GNAT1 trafficking in preventing retinal degeneration caused by phototoxic rhodopsin mutants.

## Acknowledgements

The authors thank WC Smith for the gift of the anti-GNAT1 polyclonal antibody. This research was funded by the National Science and Engineering Research Council (RGPIN-2020-05193) and the Canadian Institutes of Health Research (PJT-155937 and PJT-156072)

---

**Table 1.** Light intensity measurements across different treatments used. Light intensity measurements for the different light exposure paradigms used in Figures 4, 5 and 6. The photopic flicker involves exposure to 2.5 cd•s/m^2^ flashes of 6500K white light at 10 Hz over a 25 cd/m2, 6400K white background.

|  | Incubator, 10 min | Full ERG Protocol | Photopic Flicker |
| --- | --- | --- | --- |
| **Integrated Intensity (cd/m<sup>2</sup>)** | 169 942 cd/m<sup>2</sup> | 3495 cd/m<sup>2</sup> | 1525 cd/m<sup>2</sup> |
| **Intensity (cd/s/m<sup>2</sup>)** | 283 cd/s/m<sup>2</sup> | 5.825 cd/s/m<sup>2</sup> | 69.318 cd/s/m<sup>2</sup> |

